# Identification of meiotic recombination through gamete genome reconstruction using whole genome linked-reads

**DOI:** 10.1101/363341

**Authors:** Peng Xu, Human Genome Structural Variation Consortium, Zechen Chong

## Abstract

Meiotic recombination (MR), which transmits exchanged genetic materials between homologous chromosomes to offspring, plays a crucial role in shaping genomic diversity in eukaryotic organisms. In humans, thousands of meiotic recombination hotspots have been mapped by population genetics approaches. However, direct identification of MR events for individuals is still challenging due to the difficulty in resolving the haplotypes of homologous chromosomes and reconstructing the gamete genome. Whole genome linked-read sequencing (lrWGS) can generate haplotype sequences of mega-base pairs (N50 ~2.5Mb) after computational phasing. However, the haplotype information is still isolated in a large number of fragmented genomic regions and limited by switch errors, impeding its further application in the chromosome-scale analysis. In this study, we developed a tool MRLR (<u>M</u>eiotic <u>R</u>ecombination identification by <u>L</u>inked-<u>R</u>ead sequencing) for the analysis of individual MR events. By leveraging trio pedigree information with lrWGS haplotypes, our pipeline is sufficient to reconstruct the whole human gamete genome with 99.8% haplotyping accuracy. By analyzing the haplotype exchange between homologous chromosomes, MRLR identified 462 high-resolution MR events in 6 human trio samples from the Genome In A Bottle (GIAB) and the Human Genome Structural Variation Consortium (HGSVC). In three datasets of the HGSVC, our results recapitulated 149 (92%) previously identified high-confident MR events and discovered 85 novel events. About half (40) of the new events are supported by single-cell template strand sequencing (Strand-seq) results. We found that 332 (71.9%) MR events co-localize with recombination hotspots (>10 cM/Mb) in human populations, and MR breakpoint regions are enriched in PRDM9 and DMC1 binding sites. In addition, 48% (221) breakpoint regions were detected inside a gene, indicating these MRs can directly affect the haplotype diversity of genic regions. Taken together, our approach provides new opportunities in the haplotype-based genomic analysis of individual meiotic recombination. The MRLR software is implemented in Perl and is freely available at https://github.com/ChongLab/MRLR.

## Introduction

Homologous recombination is a molecular process in which nucleotide sequences are exchanged between similar DNA fragments (San Filippo et al. 2008). During meiosis, meiotic recombination (MR) events occur between homologous chromosomes of paternal and maternal origin, leading to novel combinations of genetic materials transmitted to gametes. This process contributes to genetic exchange across generations and provides genetic diversity for natural selections (Coop and Przeworski 2007). Investigation of MR events in individuals is important to track the law of genetic inheritance in human pedigrees (Reich et al. 2002) and illustrate the functional impact of MR on genomic variations (Jeffreys et al. 2001; Norman et al. 2009). Failure to maintain correct MR may lead to severe genetic diseases such as Down Syndrome and cancers in humans (Purandare and Patel 1997; Hassold and Hunt 2001).

MR occurrence is regulated by the molecular system of DNA double-strand break (DSB) repair. During meiosis, programmed DSBs are initiated by SPO11 via a topoisomerase-like reaction (Keeney et al. 1997). This process is proposed to be guided by the histone-lysine N-methyltransferase PRDM9 protein (Baudat et al. 2013), which contains zinc finger DNA binding motif and triggers local chromatin modification (Baudat et al. 2010; Baker et al. 2014; Pratto et al. 2014). Following DSB initiation, recombinase DMC1 and RAD51 are involved in repair of DSBs. They interact with single-stranded DNA and mediate the homologous DNA pairing and strand exchange reaction (Sehorn et al. 2004). Depending on different molecular mechanisms, DSB repair can give rise to genetic crossovers or non-crossovers (de Massy 2013). Crossovers are formed through the resolution of a double Holliday junction intermediate in which a reciprocal genetic exchange occurs between homologous chromosomes. In non-crossovers, gene conversion usually occurs via the synthesis-dependent strand annealing (SDSA) pathway, generating a unidirectional exchange of short nucleotide sequences (McMahill et al. 2007). As crossovers and non-crossovers are generated from distinct mechanisms, this study focused on large-scale MR events that are derived from crossovers.

MR events can be characterized on the scale of a population or an individual. Constructing chromosome haplotypes in which nucleotide variations are concurrently inherited is the key for MR event identification. Population genomic methods infer haplotype locally through exploiting strong correlations of single nucleotide variants (SNVs) within linkage disequilibrium (LD) blocks (Browning and Browning 2011). In human populations, about 33,000 meiotic recombination hotspots were discovered with elevated recombination activity (International HapMap et al. 2007). These hotspots are tightly clustered in 1-2 kb regions along chromosomes. In the individual human genome, MR events were studied by single-cell sequencing of gamete cells (Lu et al. 2012; Hou et al. 2013). Through sequencing a pool of gamete cells, these studies tried to reconstruct parental chromosome haplotypes, and infer meiotic crossovers for each gamete cell. With a focus on a collection of gamete cells, however, these studies are not feasible to identify inherited MR events during which only a specific gamete cell is fertilized and transmitted. To identify inherited MR events, the gamete genome needs to be reconstructed and compared with the haplotypes of parental homologous chromosomes.

Resolving a chromosome haplotype, also termed as phasing, is still challenging in human genomic studies. Several experimental approaches were developed to achieve a long range of chromosomal phasing (Ma et al. 2010; Selvaraj et al. 2013). Among them, single-cell DNA template strand sequencing (Strand-seq) has been demonstrated as an efficient approach to phase diploid genomes along the entire chromosomes (Falconer et al. 2012; Porubsky et al. 2016). In Strand-seq, a mitotic division of a parental cell is required to remove newly synthesized strands and selectively retain parental template strands in a daughter cell. Then directional sequencing library is constructed to distinguish single DNA molecules of the parental template strands (Sanders et al. 2017). Through sequencing 100 single cell libraries, Strand-seq is able to phase 60%-70% genomic SNVs (Porubsky et al. 2016). After phasing the genomes of trio samples separately, Strand-seq compares the parental and child’s haplotypes and identifies inherited MR events. As the sequenced genome is not PCR-amplified during library construction, Strand-seq has several limitations such as low coverage of phased SNVs, high switch errors for SNV phasing, and time-consuming for experimental work (Porubsky et al. 2017). In practice, Strand-seq can be combined with PacBio long read sequencing technology to achieve a high quality of phased chromosomes (Porubsky et al. 2017).

The whole genome linked-read sequencing (lrWGS) of 10X Genomics is another technical advancement to generate haplotype sequences with a relatively long range. Linked-read sequencing mainly relies on droplet-based reagent delivery system that partitions genomic DNA (gDNA) molecules with ten to several hundred kilobases into gel beads (Zheng et al. 2016; Bell et al. 2017). Then oligonucleotides with a common barcode for each partition are delivered. After high-throughput sequencing with the Illumina platform, linked-read sequencing can generate millions of short reads with unique barcodes. By anchoring the barcoded reads that are uniquely mapped to the reference, it can distinguish sequences in repetitive sequences and improve the overall alignment accuracy. On the other hand, long-range barcode information can be used to discriminate haplotypes of homologous chromosomes. In human genomes, 10X Genomics lrWGS can phase over 98% of SNVs with an N50 of ~2.5 Mb for phased blocks (Zheng et al. 2016). The phasing switch error of 10X Genomics linked-reads is about 0.025%, much lower than Pacbio (0.13%) and Strand-seq (0.32%) (Porubsky et al. 2017). However, as distant heterozygous sites lacking shared barcode information could not be connected, the linked-read sequencing method usually produces thousands of haplotype blocks fragmented in the genome.

To comprehensively and accurately identify inherited MR events, we developed a method MRLR (<u>M</u>eiotic <u>R</u>ecombination identification by <u>L</u>inked-<u>R</u>ead sequencing) which harnesses the long-range property of linked-reads and the pedigree information.

MRLR overcomes the limitation of fragmented haplotypes generated lrWGS and reconstructs the gamete genome at chromosome level with high accuracy. MRLR also showed high performance in identifying MR events in diverse human population backgrounds. Analysis of MR breakpoint regions reveals featured patterns of MR occurrence in individuals and their functional impact on shaping genetic diversity. We expect our approach can be widely applied in future studies to better understand human inheritance and disease.

## Results

### MRLR pipeline for MR event identification

MR transmits the recombined chromosomes from parents to child during gamete (sperm and egg) cell formation. To identify MR breakpoints, MRLR compares the gamete genome with the parental haplotypes (**Fig. 1A**). The pipeline consists of two procedures. The first part is to reconstruct the gamete genome. As the transmitted gamete cells already hybridized into the zygote, MRLR uses trio samples to reconstruct the gamete genome. This process is achieved by allocating fragmented haplotype blocks of the child to haplotypes of parents to build a consecutive and correct haplotype (**Fig. 1B**). As switch errors in haplotype blocks may inhibit haplotype allocation to parents, MRLR splits the phased blocks of the child into short (10 kb) bins to test haplotype relationships of each bin between child and each parent (**Fig. 1B** and **Methods**). Then, the continuous split bins will be merged if their haplotype configuration can be resolved by the haplotypes of parents. Haplotype blocks that cannot be assigned to parents will be discarded in this process. In the end, MRLR concatenates fragmented phased blocks of the child into one complete gamete haplotype derived from the paternal or maternal origin (**Fig. 2A**).

**Fig. 1.**
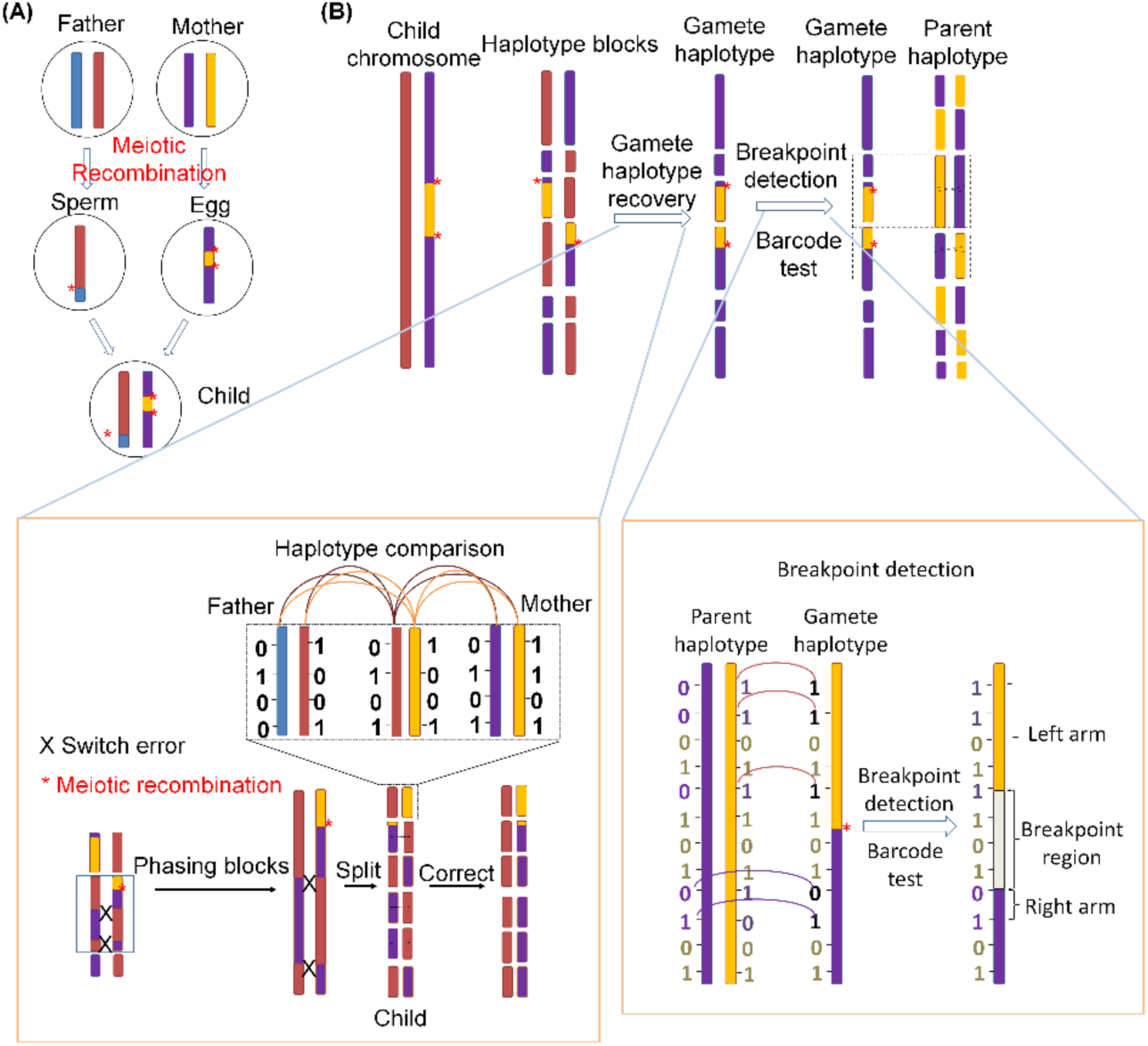
MRLR pipeline to reconstruct gamete haplotype and identify meiotic recombination events. (A) Schematic diagram of meiotic recombination. Homologous chromosomes are recombined during the meiosis process in parents and transmitted to the gametes. (B) Workflow of the pipeline. The main scheme is to infer the gamete genome through comparing parents and child haplotypes and then identify the breakpoints using the gamete and the parent haplotype information. The blue, orange, purple and yellow colors indicate different chromosome segments inherited from parents. The red asterisk indicates the meiotic recombination breakpoints. The black cross indicates switch errors in phasing. The number 0 and 1 in each haplotype indicate the reference allele and variant allele.

**Fig. 2.**
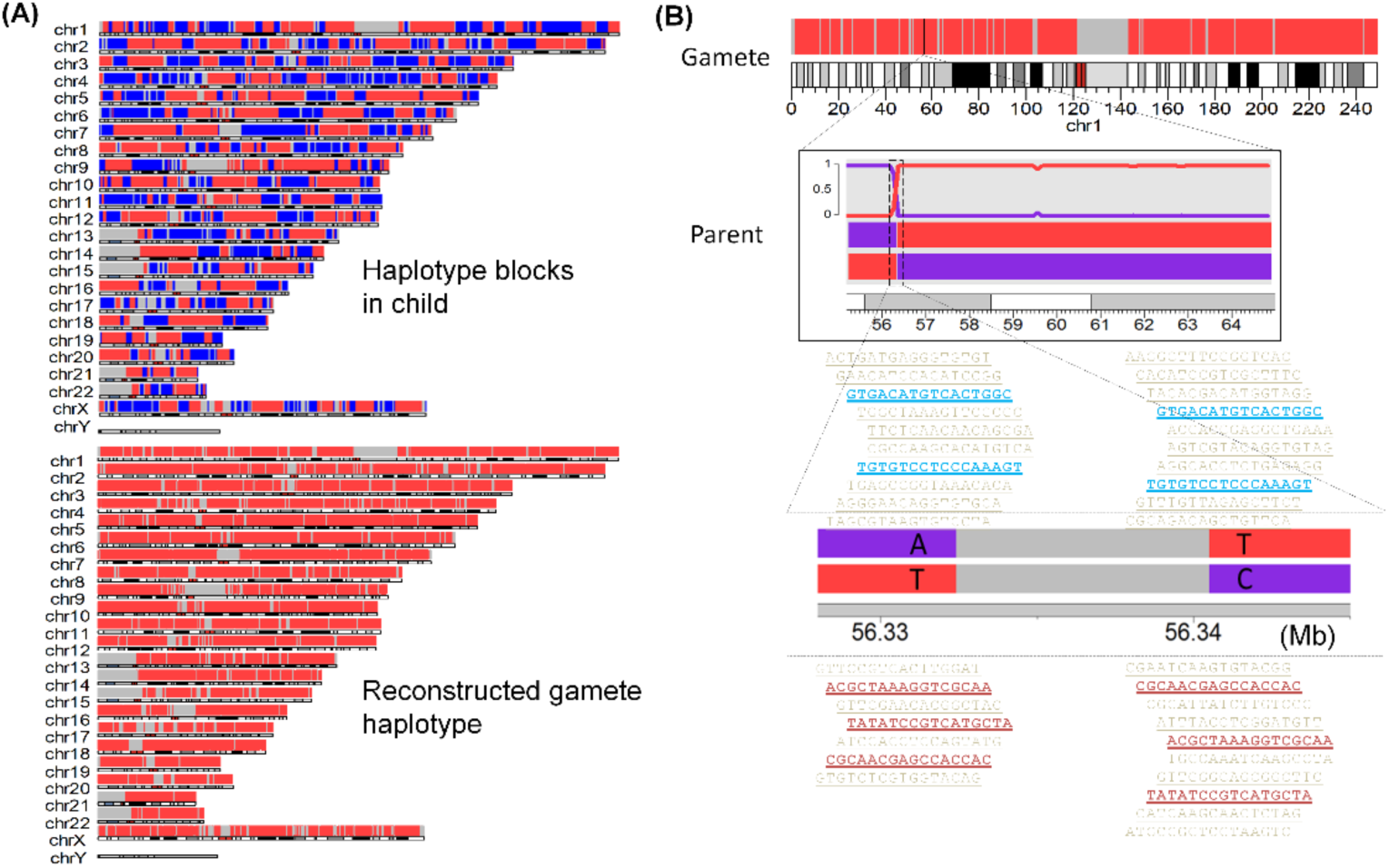
Illustration of gamete haplotype reconstruction and meiotic recombination identification. (A) gamete haplotype reconstruction from the NA12878 sample. The top part shows the original LongRanger results. The red color indicates one haplotype of the chromosome, the blue color indicates the reversed haplotype, and gray color indicates lack of phasing information. The bottom part shows the reconstructed gamete haplotype inherited from the father. (B) A meiotic recombination breakpoint identified in NA12878 father sample. The red color indicates the gamete haplotype. The line plot in the box shows the inheritance ratio of each of the father haplotypes. The red region from the father genome indicates the inherited chromosome fragments. The lower part shows the barcode information that supports the phasing of the father haplotypes. Up to ten barcodes are shown for each allele.

The second part of MRLR is to identify breakpoints of MR events in each parent. Here, MRLR uses the layout of heterozygous SNVs in parental haplotype blocks to compare their haplotype conformation with the reconstructed gamete genome (**Fig. 1B and 2B**). In a phased block of a parent, an MR event will be identified if adjacent parts of the gamete genome are assigned to two different haplotypes in the parent. Then, by a barcode testing method (**Methods** and **Supplemental Fig. S1**), MRLR reanalyzes the linked-reads in both the parent and the child that cover the MR breakpoint region. The barcode testing further validates the MR event and ensures a high-confidence call. For the MR breakpoints passed the barcode test, MRLR gathers the information of breakpoint region that spans two nearest heterozygous SNVs, as well as the flanking arms that are involved in MR events, providing the highest resolution for MRs.

### Evaluation of MRLR for MR identification

To evaluate the performance of MRLR for MR identification, we analyzed six trio lrWGS samples from the Genome In A Bottle (GIAB) (Zook et al. 2016) and the Human Genome Structural Variation Consortium (HGSVC) (Chaisson et al. 2017).

First, we evaluated the process of gamete reconstruction. To benchmark the performance of the process, we analyzed the genome NA12878 from GIAB. NA12878 is the child sample from a GAIB trio dataset and its genome has been extensively studied by multiple sequencing platforms. The reconstructed gamete genome is expected to be derived from one haplotype of the child genome. In NA12878, about 2.17 million heterozygous SNVs are phased. Among them, 2.09 million (96.5%) are located in phased regions of the reconstructed gamete genomes and 1.78 million (82.3%) SNV sites are captured in the reconstructed gamete genome. Among those captured heterozygous SNVs, 99.8% are uniformly located in one haplotype of the child genome, suggesting the gamete genomes are correctly reconstructed by MRLR (**Fig. 2A** and **Supplemental Table S1**). Among uncaptured SNVs, the majority (92.1%) are either with low sequencing quality or not reliably phased in at least one of the trio samples.

To evaluate the performance of MRLR for MR breakpoint identification, we analyzed three trio samples of Han Chinese, Puerto Rican and Yoruban Nigerian from HGSVC. These three trio samples represent different population backgrounds and a total of 162 MR events have been identified by MR analysis using a combination of Strand-seq and Pacbio method (Chaisson et al. 2017). Although without experimental validation, the MR events identified by both methods were considered as high confident events and used as ground truth in this study to estimate optimal parameters of MRLR. These parameters include the size of phased blocks occurring MR events, the number of heterozygous SNVs supporting each breakpoint arm, the number of barcodes supporting the phased breakpoint regions, the length of each breakpoint arm, and the length of breakpoint regions. Interestingly, in the Yoruban Nigerian samples, the estimated sensitivity for MR detection is always higher than 95% under a wide range of parameter settings (**Fig. 3**). In contrast, the Han Chinese and Puerto Rican are more sensitive to parameter changes. Their estimated sensitivities will drop dramatically (< 90%) if more heterozygous SNVs (> 20) and longer arm length (> 30kb) are required to define a breakpoint region (**Fig. 3B** and **3D**). The block size and supporting barcode number also affect the sensitivities when more stringent criteria are applied (**Fig. 3A** and **3C**). This suggests that the parameters of MRLR for MR breakpoint detection should be customized according to different human genetic backgrounds. The Yoruban Nigerian is from African populations, which have greater levels of genetic diversity (Campbell and Tishkoff 2008). The high diversity can facilitate phasing and thus can generate high-quality haplotypes. Consistently, based on the Long Ranger results, we found that the Yoruban Nigerian samples have longer phased lengths and less phased block numbers compared with the other two samples (**Supplemental Fig. S2** and **Supplemental Table S2**). This explains why MRLR in the Yoruban Nigerian samples is robust under different parameter settings. For the Han Chinese and the Puerto Rican samples, each parameter was optimized at the threshold that allows at least 90% detection sensitivity.

**Fig. 3.**
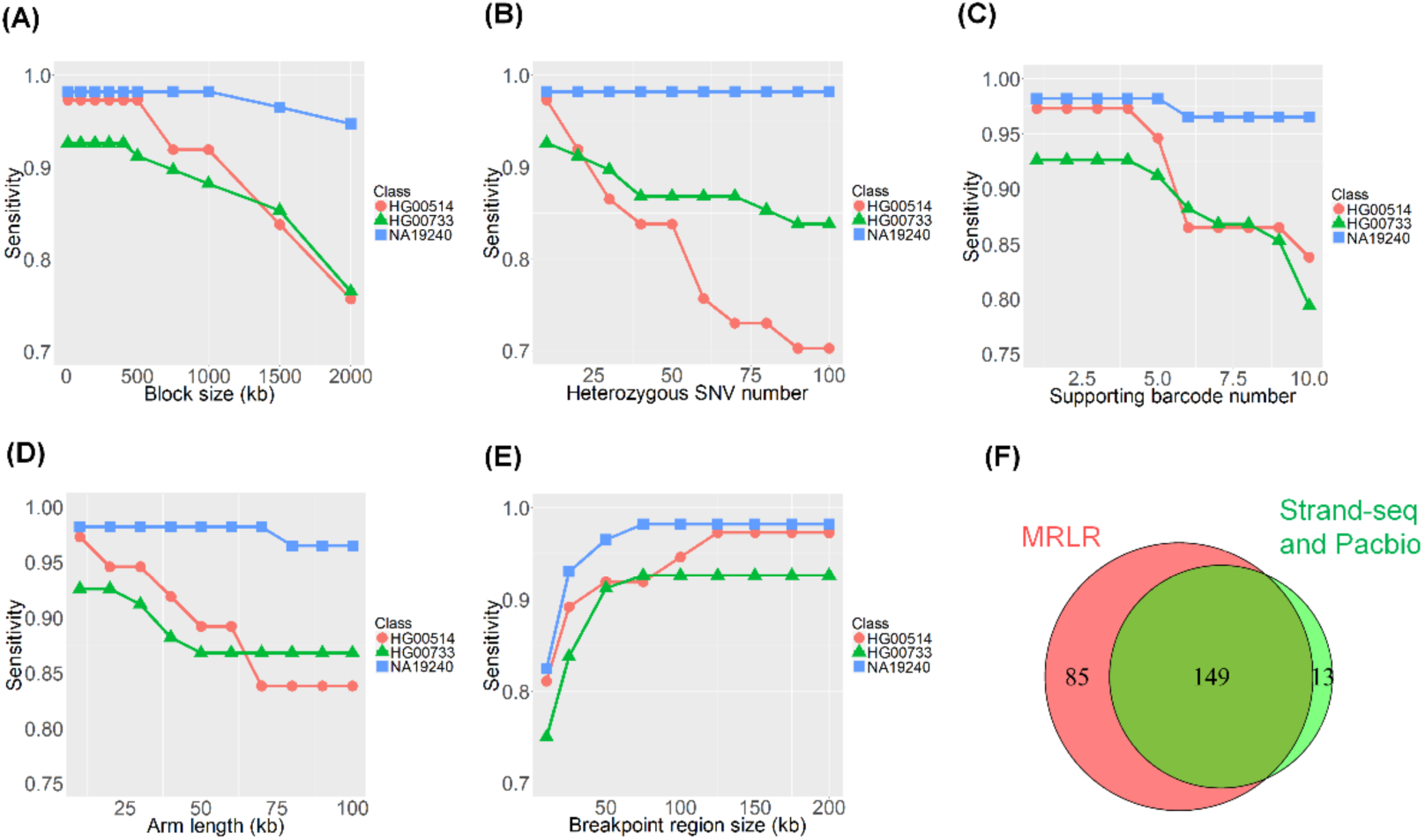
Identification of meiotic recombination events in three population groups. Different parameters including block size (A), SNV number (B), barcode number (C), arm length (D), and breakpoint region length (E) were tested in Han Chinese (HG00514), Puerto Rican (HG00733) and Yoruban Nigerian (NA19240) trio samples. In each panel, the horizontal axis represents each parameter as a cut-off to identify meiotic recombination events. The vertical axis represents detection ratios of high-confidence meiotic recombination events reported from HGSVC. (F) Venn diagram shows the overlapping MR events of HGSVC and identified by MRLR under parameters: block size >= 500kb, SNV number >= 20, barcode number >= 4, arm length >= 20 kb, and breakpoint region length <= 100kb.

By applying the optimized parameters (block size >= 500kb, SNV >= 20, barcode >= 4, arm length >= 20kb and breakpoint region <= 100 kb), MRLR totally identified 234 MR events in the three HGSVC trio samples (**Fig. 3F** and **Supplemental Table S3**). This covers 149 (92%) of high confident MR events reported by both Strand-seq and Pacbio methods (**Supplemental Table S4**). The missed 13 MR events from HGSVC are located either on the boundary or outside of the haplotype blocks, which hinders the reliable inference of MRs (**Supplemental Table S5**). Apart from high-confidence MRs of HGSVC, MRLR identified 85 new MR events. Among them, 40 new events are supported by Strand-seq results. And 31 new events have low SNV coverage (< 20 SNVs) at the flanking arms in Strand-seq results, representing the majority (69%) of MR events missed by Strand-seq (**Supplemental Table S6**). Then we compared the breakpoint resolution identified by MRLR with the Strand-seq method. The median breakpoint length of Strand-seq is about 24 kb. In contrast, the resolution of MRLR can achieve < 7 kb median length for MR breakpoint regions (**Supplemental Fig. S3**). These results suggest MRLR has high sensitivity and resolution to identify MR breakpoints.

### Features of MR breakpoints in six trio samples

Previous population genetic studies revealed MR tends to occur in 1-2 kb recombination hotspots in human genomes (International HapMap et al. 2007). However, the pattern of MR occurrence in individuals has not been systematically investigated. Thus, we applied MRLR to detect individual MR events and to analyze the features of MR breakpoints. We totally identified 462 recombination events in the six trio samples of GIAB and HGSVC (**Supplemental Table S3**). The identified MR breakpoints range from 20 to 58 events in each parent of the trio samples. The Han Chinese (HG00514) and the Yoruban Nigerian (NA19240) samples represent the populations with the lowest and highest MR events, respectively. Although 37% (169) MR events are loosely clustered with one another MR event within 1 Mb region on the chromosome, distribution of MR breakpoints regions seems to be random along the chromosomes (**Fig. 4A**). To reveal a detailed distribution of MR breakpoints on chromosomes, we zoomed in two breakpoint regions from NA24631 and NA19240 trio samples (**Fig. 4B**). These two events are closely located on chromosome 1 within a 0.5 Mb genomic region. When overlapping with the MR hotspots in human populations (International HapMap et al. 2007), we found each breakpoint region is perfectly located in hotspot regions with high recombination rates (> 40 cM/Mb) (**Fig. 4B**). This suggests the occurrence of the two breakpoint regions in individuals actually follows the rule of recombination rate in human populations. To systematically investigate the relationship between breakpoint regions and recombination rate of human populations, we extracted 50 kb sequence flanking the center of the discovered MR breakpoints and calculated the maximum recombination rate in the region. We found that 332 (71.9%) breakpoint regions overlap with recombination hotspots (> 10 cM/Mb). The permutation test showed recombination rate in MR breakpoint regions is significantly higher than randomly sampled genomic regions (p-value < 0.01) (**Supplemental Fig. S4**).

**Fig. 4.**
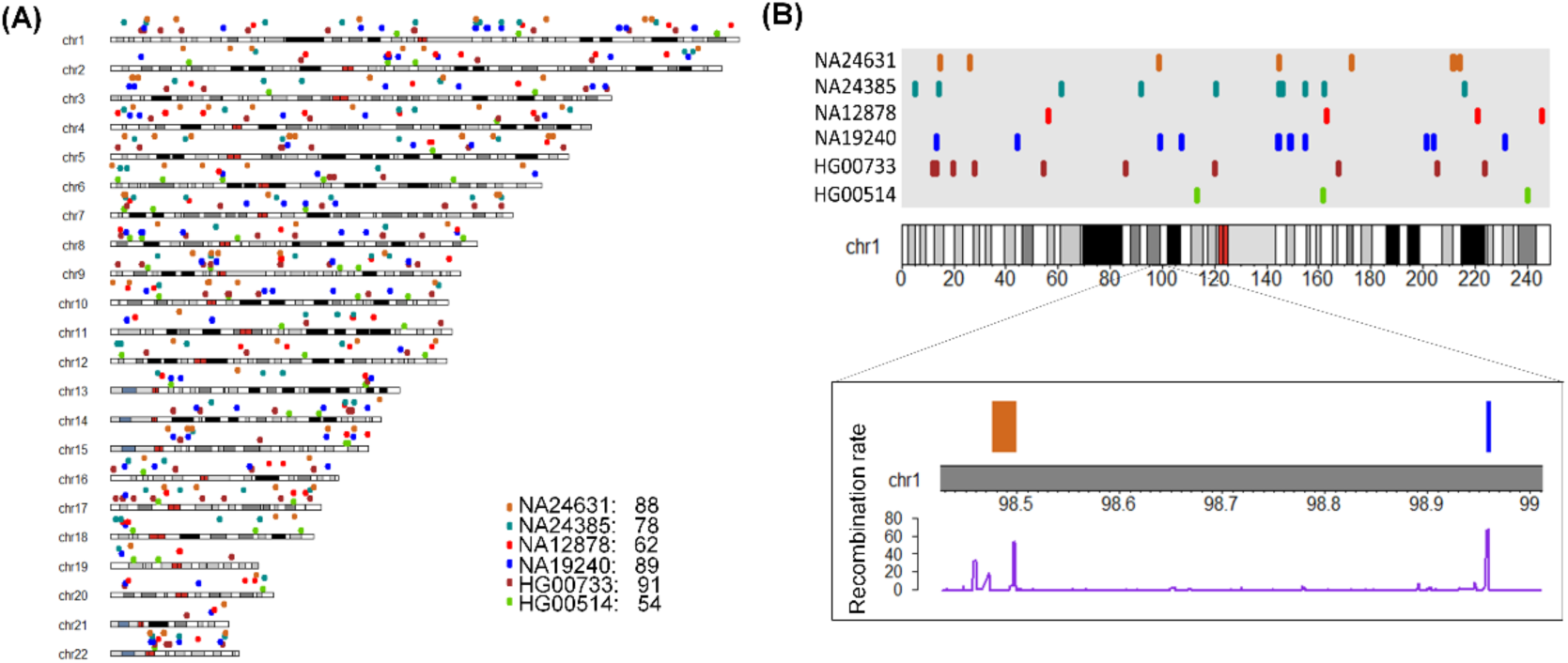
Overview of meiotic recombination breakpoints in six trio samples. (A) Distribution of MR events along the chromosomes. Each dot indicates a breakpoint region in one trio sample. (B) Two examples of breakpoint regions in chromosome 1. Up panel shows the distribution of MR events in chromosome 1. The enlarged region shows two breakpoint regions overlapping with high recombination rates in human populations. Recombination rate is obtained from HapMap Phase II.

In the six trio samples, MRLR narrowed down 74 MR breakpoints to less than 1 kb regions with the nearest heterozygous SNVs. The high resolution of breakpoint regions allows us to investigate recombination rate and functional cis-elements on a precise scale. Among breakpoint regions with 1 kb resolution, 57 MR events overlapped with the HapMap2 dataset. We extracted 500bp sequence around the center of breakpoint regions and found 42 (73.7%) sequences contain recombination hotspots in human populations (>10 cM/Mb). The breakpoint regions with 1 kb resolution have a mean recombination rate of ~30 cM/Mb. In contrast, the permutation test showed recombination rate in randomly sampled genomic regions is < 2 cM/Mb (**Fig. 5A**). This suggests the MR breakpoint regions with 1 kb resolution contain strong signals for recombination hotspot and MR occurrence.

**Fig. 5.**
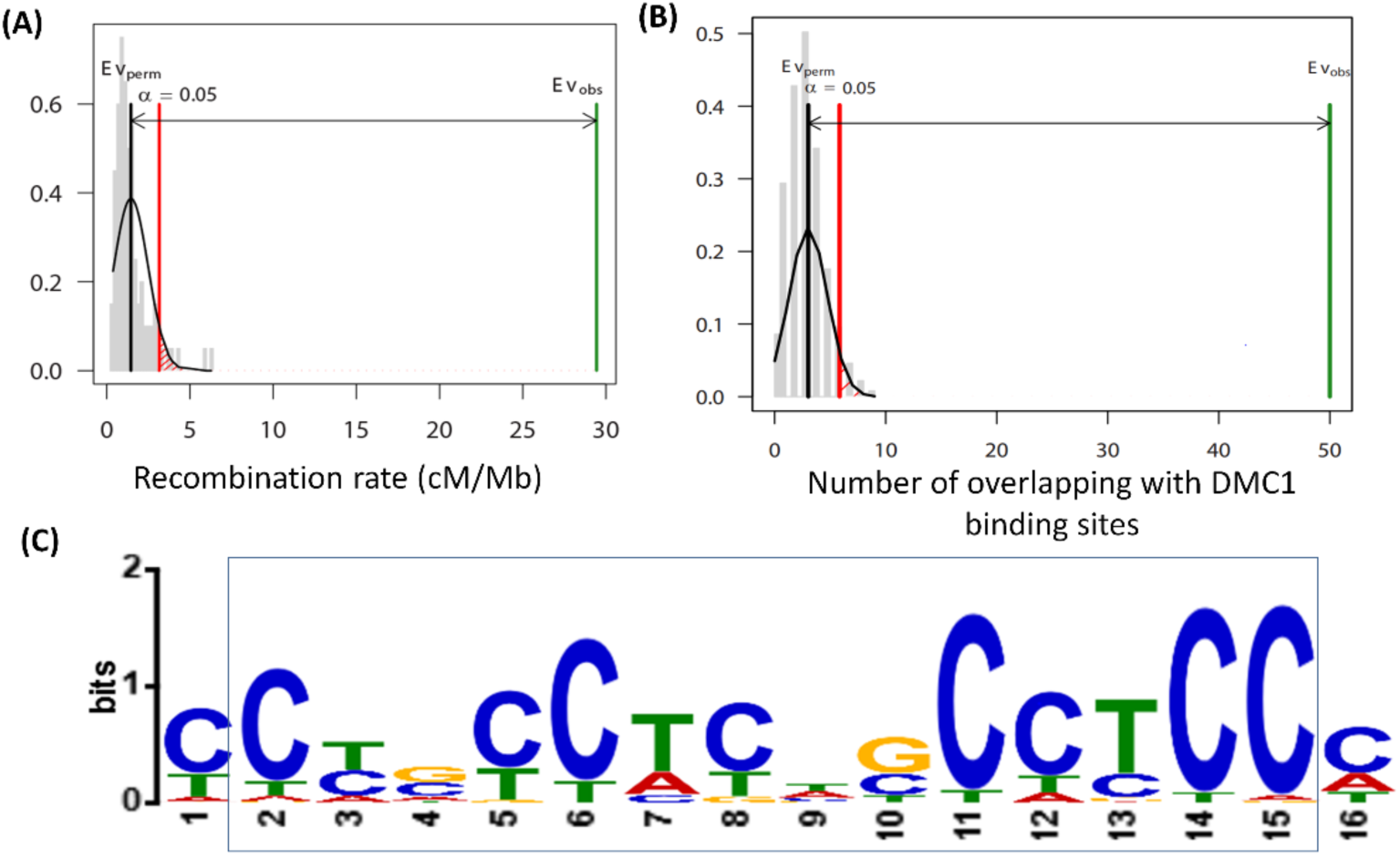
Features of DNA sequence in high resolution MR breakpoint regions. (A) Recombination rate compared between MR breakpoint regions and randomly generated sequences. (B) Overlapping of DMC1 binding sites in MR breakpoint regions and randomly generated sequences. In (A) and (B), breakpoint regions (< 1kb) in autosomes were used for analysis. The black line indicates the distribution of permutation test results. Green bar Evobs indicates the result of MR breakpoint regions. (C) The most significantly enriched DNA motif in the breakpoint regions. The boxed region shows the motif sequence similar with the binding motif of the PRDM9 protein.

DMC1 is a meiosis-specific recombinase that mediates strand exchange for DSB repair and recombination reaction (Sehorn et al. 2004; Pratto et al. 2014). Therefore, we explored whether MR breakpoint regions are enriched with DMC1 binding sites. For breakpoint regions with 1 kb resolution, 49 out of 74 (66%) breakpoint regions overlap with DMC1 binding sites. The permutation tests showed DMC1 binding sites are significantly enriched in the MR breakpoint regions (p-value < 0.01) (**Fig. 5B**). Then we used MEME toolkit to detect cis-elements that are significantly enriched in breakpoint regions with 1 kb resolution. Among all cis-elements identified, the most significantly enriched motif contains the consensus sequence C(C/T)nCCTCn(G/C)CCTCC (**Fig. 5C** and **Supplemental Fig. S5**). Interestingly, it resembles the motif (CCnCCnnnnCCnCC) that was previously identified in recombination hotspots and recognized by PRDM9 proteins for meiotic DSB initiation (Myers et al. 2008; Baudat et al. 2010; Pratto et al. 2014). Remarkably, this motif is present in all the MR breakpoint regions (< 1kb) analyzed (**Supplemental Table S7**). This implies the importance of PRDM9 proteins in guiding the MR event initiation.

### Influence of MR events on gene structure

Previous studies showed that MR is an important mechanism to increase sequence diversity for encoding genes (Jeffreys et al. 2001; Norman et al. 2009). Therefore, we investigated whether MR occurrence in individuals can directly affect gene structure and change gene haplotype. Among the 462 breakpoint regions, 221 (47.8%) breakpoint regions locate inside a gene (**Fig. 6A** and **Supplemental Table S8**). The MR containing genes include 157 protein-coding genes, 28 lincRNA genes, 14 antisense RNA genes and 13 pseudogenes (**Fig. 6B**). This suggests that about half of MR events can affect the haplotypes of genic regions. We further analyzed a breakpoint occurring on chromosome 1 of NA24631 father sample. This MR event is present in the NFIA gene which encodes a nuclear factor for transcriptional regulations and is associated with central nervous system diseases (Lu et al. 2007). The MR event occurs in the second intron of the NFIA gene (**Fig. 6C**), which overlaps with MR hotspots in human populations. Interestingly, structural variations, such as chromosome translocations and 1p31p32 microdeletion, are also located in this region (Coci et al. 2016). The intron located MR event can cause haplotype exchange of flanking exons and affect the annotated transcripts of NFIA gene. This suggests MR events can directly impact gene structure and increase haplotype diversity.

**Fig. 6.**
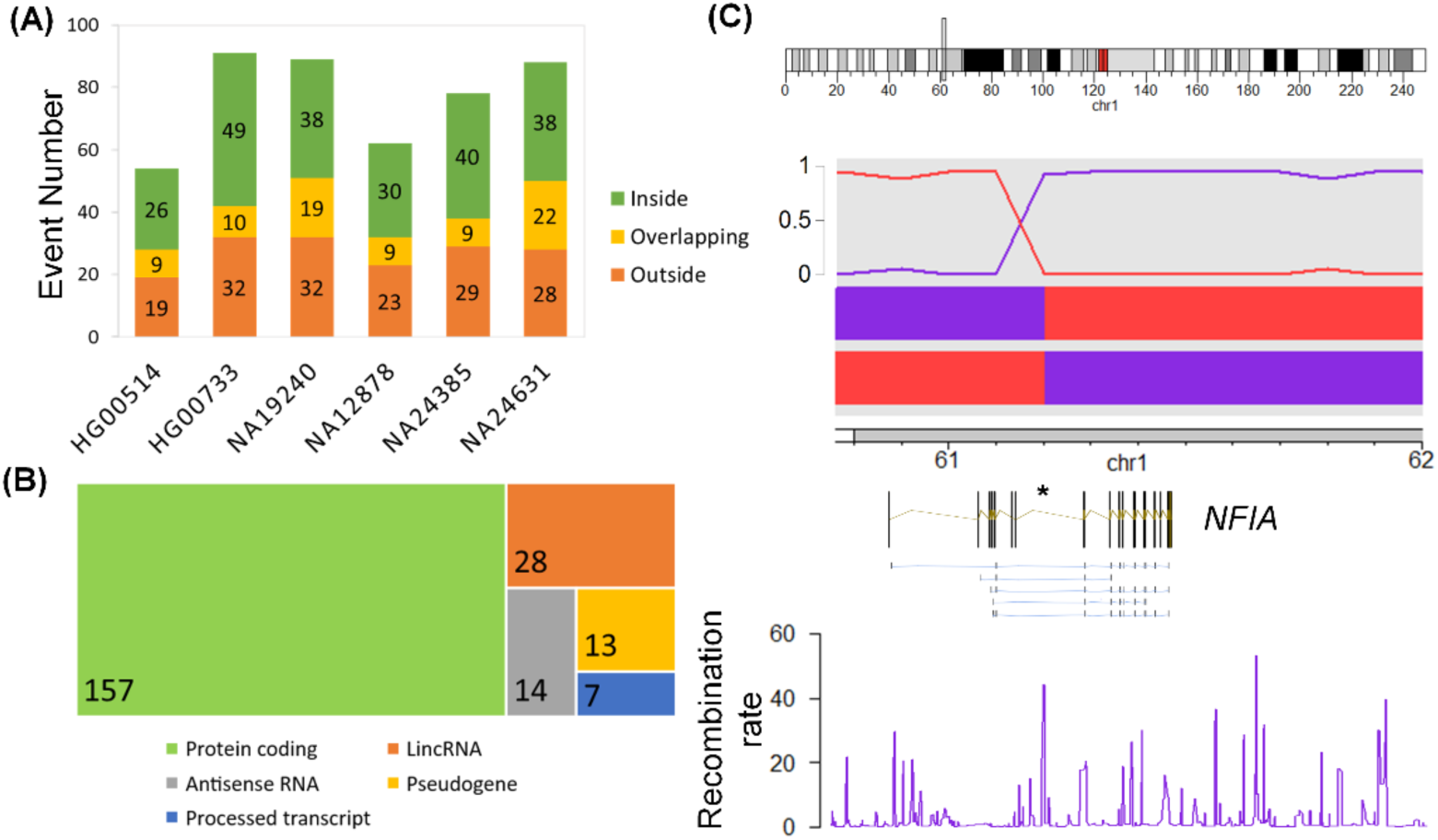
The occurrence of meiotic recombination events in genic regions. (A) The MR occurrence with respect to gene regions. (B) The type of genes that contain a meiotic recombination event. (C) An example of an MR event that occurs inside of the *NFIA* gene. From top to bottom shows the chromosomal location, inheritance ratio of two parental haplotypes, *NFIA* gene structure, *NFIA* transcripts, and recombination rate in the region, respectively.

## Discussion

In humans, recombination in meiosis is a fundamental process to promote genetic material exchange. Identifying and characterizing MR events in individual families are important to understand genetic inheritance across generations and investigate haplotype-based gene interactions in human diseases. In this study, we developed a pipeline, MRLR, to reconstruct the gamete genome and identify MR events in trio families based on the lrWGS platform. Compared with Strand-seq which relies on single-cell sequencing method, lrWGS uses the Illumina sequencing platform which is capable to produce dense and accurate SNVs. Consistent with this, our reconstructed human gamete genome can capture 82.3% of heterozygous SNVs conveyed to the offspring and achieve 99.8% phasing accuracy. This ratio is higher than Strand-seq method, which can only phase 60% - 70% genomic SNVs with 99.3% concordance in 100 single cell libraries (Porubsky et al. 2016). The rich information of phased SNVs also assists to refine MR regions to precise locations. Compared with Strand-seq, MRLR can narrow down MR breakpoint length to a more precise scale (7kb vs 24kb). We also discovered new MR events that were missed in Strand-seq results (**Supplemental Table S6**). These results showed high performance of our method to analyze meiotic recombination in human genomes.

Although MR patterns have been described in population genetic studies, our pipeline allows detailed characterization of MR events in individuals. Using MR breakpoint regions with 1kb resolution, our results revealed PRDM9 and DMC1 binding sites are significantly enriched in MR breakpoint regions. Remarkably, we observed PRDM9 binding motif is present in all of the high-resolution MR breakpoint regions. This emphasizes the importance of PRDM9 guidance in the regulation of MR occurrence. PRDM9 binding sites are polymorphic and largely overlap with repetitive sequences such as Alu elements (Myers et al. 2008). These polymorphic binding sites make MR occurrence dynamic and specific in different individuals. Consistent with this, about 70% MR breakpoint regions overlapping with recombination hotspots (> 10 cM/Mb) of human populations, suggesting a portion of MR breakpoints occur in the low frequency of populations, or even private to certain individuals. Therefore, when applied to individual families, our pipeline may facilitate future work to construct a high-resolution map of genetic inheritance and identify disease-related recombination events.

In addition to promoting genetic exchange between homologous chromosomes, MR was also considered as an important mechanism to increase gene diversity (Jeffreys et al. 2001; Norman et al. 2009). Our results showed about half of MR events directly occur in genic regions in different individuals. Consistent with the observation that incorrect MR events are related gene malfunctions and human diseases (Purandare and Patel 1997; Hassold and Hunt 2001), our results showed that MR events have functional impacts on shaping gene structures and haplotype diversity. Apart from MR events mediated by crossovers, gene conversions generated by non-crossovers represent another category of unidirectional exchange of short nucleotide sequences. Gene conversions may also influence gene haplotype through exchanging short regions of a few SNVs (Jeffreys and May 2004). However, local errors from SNV calling and haplotype phasing pose challenges for lrWGS to identify these events correctly. Incorporating multiple sequencing platforms may be a future direction to discover non-crossover related recombination events.

To summarize, our study presents a lrWGS based pipeline to achieve gamete haplotype reconstruction and MR events identification in an individual human family. The dense haplotype and SNV information revealed by our method will facilitate detailed characterization of human genetic inheritance and interactions in future studies. As homologous recombination and DSB repair pathways are closely related, the MR analysis will also gain insights into human diseases in the personalized medicine era.

## Methods

### 10x linked-read sequencing data retrieval and pre-processing

The six trio samples for linked-read sequencing analysis were obtained from two public resources. Three trio datasets were from the Genome in a Bottle Consortium (GIAB). They include Han Chinese (Son NA24631, Mother NA24695, Father NA24694), Ashkenazim Jewish (Son NA24385, Mother NA24143, Father NA24149) and Caucasian (Daughter NA12878, Mother NA12892, Father NA12891). Three trio datasets were from Human Genome Structural Variation Consortium (HGSVC). They include Han Chinese (Son HG00514, Mother HG00513, Father HG00512), Puerto Rican (Son HG00733, Mother HG00732, Father HG00731) and Yoruban Nigerian (Son NA19240, Mother NA19238, Father NA19239). These samples were sequenced by the Chromium Genome Platform of 10X Genomics. The raw datasets were processed by the Long Ranger software (Zheng et al. 2016). SNVs were called by Freebayes (Garrison and Marth 2012). Samples were aligned and processed against human genome reference GRCh38.

For each trio dataset, the haplotype information was retrieved from the VCF files of Long Ranger outputs and combined into one file. Each row represents an SNV. Each column represents the SNV haplotype information in Father, Mother and Child sample. Initial filtering of SNVs was applied to each of the trio samples: 1) the SNVs should pass all the filters of Long Ranger software; 2) there are minimal 5 read support of the observed variant; 3) the average base (Phred) quality is higher than 30; 4) there is no violation of Mendelian inheritance in a trio family; 5) the called variant should be phased in a phasing block. The filtered SNVs are regarded as high-quality SNVs and used for downstream analysis.

### Gamete genome reconstruction

We reconstructed the gamete genome through haplotype comparison between parents and child. The main scheme is to use the pedigree information to distinguish the two homologous chromosomes in the child. In the child, haplotype blocks are not always identical to one of the parent haplotype due to the influence of meiotic recombination, genetic mutations and switch errors during phasing. To mitigate these factors, we split large haplotype blocks into 10kb region in the child. Then we test the relationship between the split haplotype blocks in the child and the parent haplotype. At each split haplotype block of the child, let *cp* be the nucleotide of one haplotype and *cq* be the nucleotide of the other haplotype. Let *fp* and *fq* be the nucleotides of two haplotypes in the father. At a given site *i*, the relationship score between *cp* and *fp* is:

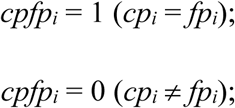

Let *n* be the total number SNVs in the split haplotype region, we calculate the relationship score ratio between *cp* and *fb*.

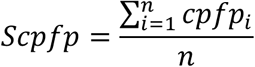

Similarly, we calculate the relationship score *Scpfq* (between cp and fq), *Scqfp* (between cq and fp), *Scqfq* (between cq and fq) in the haplotype block. Then we have a sample *X* = {*Scpfp*, *Scpfq*, *Scqfp*, *Scqfq*}. Let *X*_(*i*)_ be the *i*th order statistic of sample *X*. If *X*_(*4*)_ >0.6 and *X*_(*4*)_ - **X*_(*3*)_* > 0.1, we assign inheritance relationship to the parent-child pair in *X*_(*4*)_. We also considered the situation that when *X*_(*4*)_ and *X*_(*3*)_ involve the same haplotype in the child but they come from different haplotypes in one parent. This may be caused by phasing switch errors in the parent genome. In this case, haplotype relationship can also be assigned to the parent-child pair if *X*_(*3*)_ > 0.6, *X*_(*3*)_ - *X*_(*2*)_ > 0.1 and *X*_(*4*)_ - *X*_(*3*)_ <= 0.1.

We calculated the haplotype relationship score between the child and each parent separately and then combined the haplotype relationship from the two parents together. The combined haplotype relationship was categorized into four classes: 1) high confidence when one haplotype is assigned to father and the other haplotype is assigned to mother; 2) low confidence when only one haplotype can be assigned to a parent; 3) no evidence when no haplotype can be assigned to a parent; 4) error when two haplotypes are assigned to the same parent. To reconstruct the gamete genome, we selected the continuous haplotype blocks that are supported by at least one high confident phasing blocks without an error gap. In this process, we realign the haplotype according to their relationship to the parent to get a uniformly phased haplotype across the whole genome.

### Meiotic breakpoint region identification

The meiotic breakpoint was identified by comparing the parent haplotype and the reconstructed gamete haplotype. In each phasing block of the parent, heterozygous SNVs were used to compare with the gamete haplotype. Let *gp* be the nucleotide of the gamete inherited from father. Let *fp* and *fq* be the nucleotides of the two haplotypes in the father. At a given site *i*, the relationship score between parent and child is:

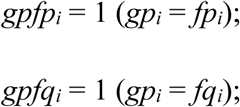

A breakpoint will be detected if its flanking region satisfies Σ*gpfp* >= 10 and Σ*gpfq* >= 10. The default flanking arm length was set >= 10 kb. The breakpoint region was defined as the nearest region of the two heterozygous SNVs that support each of the flanking arms.

We used the barcode test to ensure breakpoint regions correctly phased. It compares barcode sequence at each SNV site iteratively until the breakpoint end (**Supplemental Fig. S1**). For each SNV, we require at least 3 barcodes support the reference allele and 3 barcodes support the variant reference. Let *p* and *q* be the barcodes that support the two alleles on each of the haplotypes. Let *m* and *n* be the first and last SNV site in the breakpoint region. We pooled barcodes of three alleles from each haplotype at the boundary of left arm and made the set *PM* = {*p*_*m*-*3*_, *P*_*m*-*2*_, *p*_*m*-*1*_} and *QM* = {*q*_*m*-*3*_, *q*_*m*-*2*_, *q*_*m*-*1*_}. At SNV site *i* of the breakpoint region, we calculate the supported barcode number:

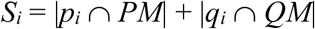

And the denied barcode number:

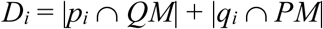

If the barcode support ratio *S_i_*/(*S_i_* + *D_i_*) >= 0.8, the SNV site is regarded as reliably phased. Then we will put the barcodes into the *PM* set {*p_*m*-*2*_*, *p*_*m*-*1*_, *p_i_*} and *QM* set {*q*_*m*-*2*_, *q*_*m*-*1*_, *q_i_*} to test the next SNV site. If the barcode support ratio *S_i_*/(*S_i_* + *D_i_*) < 0.8, this site will be discarded and the barcode sets will remain the same. The test lasts until the end of breakpoint region.

Then we pooled barcodes of three alleles from each haplotype at the boundary of the right arm and made the set *PN* = {*p*_*n*+*1*_, *p*_*n*+*2*_, *p*_*n*+*3*_} and *QN* = {*q*_*n*+*1*_, *q*_*n*+*2*_, *q*_*n*+*3*_}. We compared the barcode overlap between the left arm and right arm. The supported barcode number is:

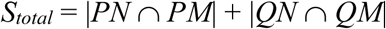

And the denied barcode number is:

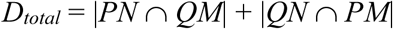

If the barcode support ratio *S_total_*/(*S_total_* + *D_total_*) >= 0.8, the breakpoint region is regarded as reliably phased. The final breakpoint regions should pass the barcode test in both in parent and child.

### Analysis of DNA sequence features in MR breakpoint regions

Recombination rate was obtained from the genetic map of HapMap Phase II. SNP coordinates in the chromosome were converted from build 37 to build 38 by UCSC liftover tool. For statistical analysis, we use RegioneR to randomly generate chromosome region with the same length of the breakpoint regions. Then we calculated the mean recombination rate in the sampled regions. We did 100 permutations to generate the distribution of recombination rates in the sampled regions. To analyze the overlapping with the DMC1 binding site, ChIP-seq results of DMC1 protein in humans were used to test the sequence enrichment (Pratto et al. 2014). RegioneR was used to generate the permutation sequences and count the overlapping between sequence regions with the DMC1 binding site. To exclude the influence of sex chromosome in estimating the recombination rate, only breakpoint regions in autosomes were used for statistical analysis.

We used breakpoint regions less than 1kb to identify enriched cis-elements. We calculate the center of the breakpoint regions and extract a 500bp sequence around the center. Then MEME software (v4.12) was used to identify cis-elements that are enriched in the 1kb sequences. The cis-element window was set between 8bp and 16bp, assuming 0 or 1 occurrence per sequence in the given strand and the reverse complement strand.

### Gene function analysis

We annotated the location of breakpoint regions through overlapping with the gene coordinates from GENCODE database (release 27). Locations of breakpoint regions were grouped into three classes based on whether they are totally inside a gene, overlapping with a gene region or totally outside a gene. For the breakpoint regions that are inside a gene, we extract the gene types based on the gtf annotation file. The information of gene-disease relationship was obtained from BioMart in Ensembl database, which was incorporated from Developmental Disorders Genotype-Phenotype Database (DDG2P), OMIM Morbid Map and Orphanet databases.

### Data access

The trio datasets used in this study can be accessed from the Genome in a Bottle Consortium (GIAB) (ftp://ftp-trace.ncbi.nlm.nih.gov/giab/ftp/release/) and Human Genome Structural Variation Consortium (HGSVC) (ftp://ftp.1000genomes.ebi.ac.uk/vol1/ftp/data_collections/hgsv_sv_discovery/).

## Acknowledgments

This work was supported by the University of Alabama at Birmingham research start-up fund to Z.C. P.X. is supported by the Institute for Precision Cardiovascular Medicine of the American Heart Association (AHA) Institutional Data Fellowship (17IF33890015).

## References

Baker CL, Walker M, Kajita S, Petkov PM, Paigen K. 2014. PRDM9 binding organizes hotspot nucleosomes and limits Holliday junction migration. Genome Res 24: 724–732.

Baudat F, Buard J, Grey C, Fledel-Alon A, Ober C, Przeworski M, Coop G, de Massy B. 2010. PRDM9 Is a Major Determinant of Meiotic Recombination Hotspots in Humans and Mice. Science 327: 836.

Baudat F, Imai Y, de Massy B. 2013. Meiotic recombination in mammals: localization and regulation. Nat Rev Genet 14: 794–806.

Bell JM, Lau BT, Greer SU, Wood-Bouwens C, Xia LC, Connolly ID, Gephart MH, Ji HP. 2017. Chromosome-scale mega-haplotypes enable digital karyotyping of cancer aneuploidy. Nucleic Acids Res 45: e162.

Browning SR, Browning BL. 2011. Haplotype phasing: existing methods and new developments. Nature Reviews Genetics 12: 703.

Campbell MC, Tishkoff SA. 2008. African genetic diversity: implications for human demographic history, modern human origins, and complex disease mapping. Annu Rev Genomics Hum Genet 9: 403–433.

Chaisson MJP, Sanders AD, Zhao X, Malhotra A, Porubsky D, Rausch T, Gardner EJ, Rodriguez O, Guo L, Collins RL et al. 2017. Multi-platform discovery of haplotype-resolved structural variation in human genomes. bioRxiv doi:10.1101/193144.

Coci EG, Koehler U, Liehr T, Stelzner A, Fink C, Langen H, Riedel J. 2016. CANPMR syndrome and chromosome 1p32-p31 deletion syndrome coexist in two related individuals affected by simultaneous haplo-insufficiency of CAMTA1 and NIFA genes. Mol Cytogenet 9: 10.

Coop G, Przeworski M. 2007. An evolutionary view of human recombination. Nature Reviews Genetics 8: 23.

de Massy B. 2013. Initiation of meiotic recombination: how and where? Conservation and specificities among eukaryotes. Annu Rev Genet 47: 563–599.

Falconer E, Hills M, Naumann U, Poon SS, Chavez EA, Sanders AD, Zhao Y, Hirst M, Lansdorp PM. 2012. DNA template strand sequencing of single-cells maps genomic rearrangements at high resolution. Nat Methods 9: 1107–1112.

Garrison E, Marth G. 2012. Haplotype-based variant detection from short-read sequencing. arXiv preprint arXiv:12073907.

Hassold T, Hunt P. 2001. To err (meiotically) is human: the genesis of human aneuploidy. Nat Rev Genet 2: 280–291.

Hou Y, Fan W, Yan L, Li R, Lian Y, Huang J, Li J, Xu L, Tang F, Xie XS et al. 2013. Genome analyses of single human oocytes. Cell 155: 1492–1506.

International HapMap C Frazer KA Ballinger DG Cox DR Hinds DA Stuve LL Gibbs RA Belmont JW Boudreau A Hardenbol P et al. 2007. A second generation human haplotype map of over 3.1 million SNPs. Nature 449: 851–861.

Jeffreys AJ, Kauppi L, Neumann R. 2001. Intensely punctate meiotic recombination in the class II region of the major histocompatibility complex. Nat Genet 29: 217–222.

Jeffreys AJ, May CA. 2004. Intense and highly localized gene conversion activity in human meiotic crossover hot spots. Nat Genet 36: 151–156.

Keeney S, Giroux CN, Kleckner N. 1997. Meiosis-specific DNA double-strand breaks are catalyzed by Spo11, a member of a widely conserved protein family. Cell 88: 375–384.

Lu S, Zong C, Fan W, Yang M, Li J, Chapman AR, Zhu P, Hu X, Xu L, Yan L et al. 2012. Probing meiotic recombination and aneuploidy of single sperm cells by whole-genome sequencing. Science 338: 1627–1630.

Lu W, Quintero-Rivera F, Fan Y, Alkuraya FS, Donovan DJ, Xi Q, Turbe-Doan A, Li QG, Campbell CG, Shanske AL et al. 2007. NFIA haploinsufficiency is associated with a CNS malformation syndrome and urinary tract defects. PLoS Genet 3: e80.

Ma L, Xiao Y, Huang H, Wang Q, Rao W, Feng Y, Zhang K, Song Q. 2010. Direct determination of molecular haplotypes by chromosome microdissection. Nat Methods 7: 299–301.

McMahill MS, Sham CW, Bishop DK. 2007. Synthesis-dependent strand annealing in meiosis. PLoS Biol 5: e299.

Myers S, Freeman C, Auton A, Donnelly P, McVean G. 2008. A common sequence motif associated with recombination hot spots and genome instability in humans. Nat Genet 40: 1124–1129.

Norman PJ, Abi-Rached L, Gendzekhadze K, Hammond JA, Moesta AK, Sharma D, Graef T, McQueen KL, Guethlein LA, Carrington CV et al. 2009. Meiotic recombination generates rich diversity in NK cell receptor genes, alleles, and haplotypes. Genome Res 19: 757–769.

Porubsky D, Garg S, Sanders AD, Korbel JO, Guryev V, Lansdorp PM, Marschall T. 2017. Dense and accurate whole-chromosome haplotyping of individual genomes. Nat Commun 8: 1293.

Porubsky D, Sanders AD, van Wietmarschen N, Falconer E, Hills M, Spierings DC, Bevova MR, Guryev V, Lansdorp PM. 2016. Direct chromosome-length haplotyping by single-cell sequencing. Genome Res 26: 1565–1574.

Pratto F, Brick K, Khil P, Smagulova F, Petukhova GV, Camerini-Otero RD. 2014. DNA recombination. Recombination initiation maps of individual human genomes. Science 346: 1256442.

Purandare SM, Patel PI. 1997. Recombination hot spots and human disease. Genome Res 7: 773–786.

Reich DE, Schaffner SF, Daly MJ, McVean G, Mullikin JC, Higgins JM, Richter DJ, Lander ES, Altshuler D. 2002. Human genome sequence variation and the influence of gene history, mutation and recombination. Nat Genet 32: 135–142.

San Filippo J, Sung P, Klein H. 2008. Mechanism of eukaryotic homologous recombination. Annu Rev Biochem 77: 229–257.

Sanders AD, Falconer E, Hills M, Spierings DCJ, Lansdorp PM. 2017. Single-cell template strand sequencing by Strand-seq enables the characterization of individual homologs. Nat Protoc 12: 1151–1176.

Sehorn MG, Sigurdsson S, Bussen W, Unger VM, Sung P. 2004. Human meiotic recombinase Dmc1 promotes ATP-dependent homologous DNA strand exchange. Nature 429: 433–437.

Selvaraj S, J RD, Bansal V, Ren B. 2013. Whole-genome haplotype reconstruction using proximity-ligation and shotgun sequencing. Nat Biotechnol 31: 1111–1118.

Zheng GX, Lau BT, Schnall-Levin M, Jarosz M, Bell JM, Hindson CM, Kyriazopoulou-Panagiotopoulou S, Masquelier DA, Merrill L, Terry JM et al. 2016. Haplotyping germline and cancer genomes with high-throughput linked-read sequencing. Nat Biotechnol 34: 303–311.

Zook JM, Catoe D, McDaniel J, Vang L, Spies N, Sidow A, Weng Z, Liu Y, Mason CE, Alexander N et al. 2016. Extensive sequencing of seven human genomes to characterize benchmark reference materials. Sci Data 3: 160025.

